# Systematic discovery of mutation-directed neo-protein-protein interactions in cancer

**DOI:** 10.1101/2021.10.03.462422

**Authors:** Xiulei Mo, Qiankun Niu, Andrey A. Ivanov, Yiu Huen Tsang, Cong Tang, Changfa Shu, Alafate Wahafu, Sean P. Doyle, Danielle Cicka, Xuan Yang, Dacheng Fan, Matthew A. Reyna, Lee A.D. Cooper, Carlos S. Moreno, Wei Zhou, Taofeek Owonikoko, Sagar Lonial, Fadlo R. Khuri, Yuhong Du, Suresh S. Ramalingam, Gordon B. Mills, Haian Fu

## Abstract

Comprehensive sequencing of patient tumors reveals numerous genomic mutations across tumor types that enable tumorigenesis and progression. A subset of oncogenic driver mutations results in neomorphic activity where the mutant protein mediates functions not engaged by the parental molecule. Here, we identify prevalent variant-enabled neomorph-protein-protein interactions (neoPPI) with a *q*uantitative *H*igh *T*hroughput *d*ifferential *S*creening (qHT-dS) platform. Coupling of highly sensitive BRET biosensors with miniaturized co-expression in an ultra-HTS format allows large-scale monitoring of interactions of wild-type and mutant variant counterparts with a library of cancer-associated proteins in live cells. Screening of 13,392 interactions with 1,474,560 data points revealed a landscape of gain-of-interactions encompassing both oncogenic and tumor suppressor mutations. For example, the recurrent BRAF V600E lesion mediates KEAP1 neoPPI, rewiring a BRAF^V600E^-KEAP1 signaling axis and creating collateral vulnerability to NQO1 substrates, offering a combination therapeutic strategy. Thus, cancer genomic alterations can create neo-interactions, informing variant-directed therapeutic approaches for precision medicine.

## INTRODUCTION

Genomic data provide a compelling structural framework to infer functional significance and to nominate potential cancer driver genes and therapeutic targets (Hahn et al., 2021; Lawrence et al., 2014; Martinez-Jimenez et al., 2020). Gene-level hotspot mutation profiling has provided valuable insights into genotypes that define clinical phenotypes and predict drug response (Garnett et al., 2012; Garraway and Lander, 2013; Vogelstein et al., 2013). However, emerging observations of context-dependent distinct impacts of mutations, beyond recurrently mutated genes, indicate their potentially diverse functional consequences and inequivalent clinical meaning (Burd et al., 2014; Chang et al., 2016; Menzies et al., 2012; Vivanco et al., 2012; Westcott et al., 2015). Therefore, translating this cancer mutation information into therapeutic applications at mutated amino acid residue resolution presents both an unprecedented challenge and opportunity for identifying lineage- and mutant allele-specific targets, biomarkers, and therapeutic vulnerabilities (Chang et al., 2016; Chin et al., 2011; Eifert and Powers, 2012; Schreiber et al., 2010).

Some mutations are found in driver genes that encode actionable targets, but the majority are located in genes without direct connection to known drugs, or in genes encoding “undruggables” such as many adaptor proteins or tumor suppressors (Ivanov et al., 2013; Vogelstein et al., 2013). These “undruggable” proteins exert their functions primarily through interactions with other cellular components through protein-protein interaction (PPI) networks (Arkin et al., 2014; Ivanov et al., 2013; Li et al., 2017). Thus, understanding how cancer driver mutations are integrated within cellular signalling networks through altered PPIs may lead to novel strategies for pathway perturbation to reach the “undrugged” space of the cancer genome (Fig. 1A).

**Figure 1.**
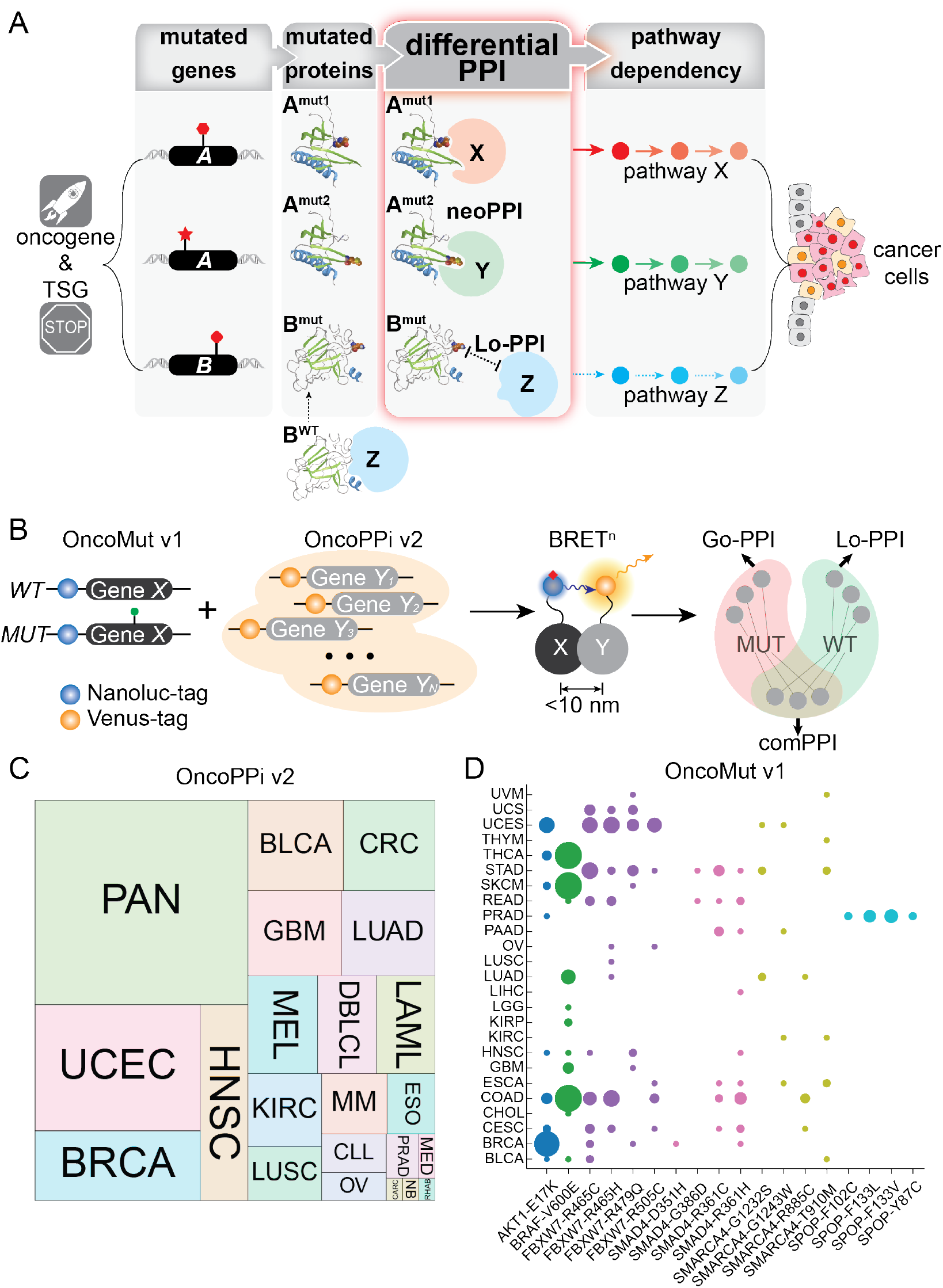
Schematic illustration of qHT-dS platform for systematic differential PPI discovery. See also Figure S1-2. **(A)** Cancer driver mutations are translated to new residues at the surface of oncoproteins, which may enhance affinities for certain proteins (X, Y) over the WT to induce oncogenic pathways while mutated residues on certain genes, such as tumor suppressors, may hinder the interaction of bound proteins (Z), rewiring the oncogenic signaling. These differential PPIs enabled by driver mutations bridge the cancer genotype to phenotype through re-wired oncogenic pathways. **(B)** Components of the qHT-dS platform. Binary parallel interaction profiling was performed by scanning NLuc-tagged WT and MUT genes in the OncoMut v1 library against Venus-tagged OncoPPi v2 library genes using the BRET^n^ detection system to identify three possible classes of PPIs, including mutation enhanced Go-PPIs, mutation-reduced Lo-PPI and common PPIs. **(C)** Square pie chart illustrating the tumor types by the distribution of genes in OncoPPi v2 library. **(D)** Bubble plot showing the tumor driver mutations, and their frequency and tumor lineage distributions in OncoMut v1 library.

Missense mutations are common genetic alterations contributing to tumorigenesis (Martinez-Jimenez et al., 2020; Vogelstein et al., 2013). The mutant (MUT) alleles may create neo-epitopes on the encoded proteins (Fig. 1A). Such mutations alter the intrinsic properties of many of the encoded proteins. It is also possible that those neo-epitopes may generate new docking sites to induce neo-interactions, or act as impediments to weaken existing interactions. Such neo-epitope-triggered differential PPIs (Df-PPI) may dictate re-wired oncogenic programs that underly the dysregulated growth and spread of tumor cells and may reveal much-needed tumor-specific molecular targets for therapeutic intervention. However, the prevalence and spectrum of neo-interactions triggered by mutations in oncogenes and tumor suppressors remains to be established. How to systematically discover those mutated residue-directed PPIs in a physiologically relevant cellular environment poses a major challenge.

To address these challenges, we have established a BRET^n^ technology-based differential PPI discovery platform that allows comparative screening of wild-type (WT) and MUT allele counterparts for detection of differential interactions with cancer-associated proteins in live mammalian cells (Fig. 1B and S1). The high throughput and quantitative nature of the technology, termed *q*uantitative *H*igh *T*hroughput *d*ifferential *S*creening (qHT-dS), enabled the systematic identification of differential WT and MUT interaction proteins for 24 alleles from the examination of 13,392 pairs of potential interactions. Analysis of high-stringency Df-PPI datasets revealed widespread gain-of-interactions induced by missense mutations of both oncogenes and tumor suppressors. Examination of mutation-directed neomorph PPIs (neoPPI) revealed potential new mechanisms and signaling pathways for well-studied mutated forms of oncogenes and tumor suppressors. The BRAF^V600E^/KEAP1 interaction was selected for validation using a panel of cellular and biochemical molecular interaction assays that advanced it to a validated neoPPI status. This neoPPI redirects BRAF^V600E^ to upregulate NRF2-mediated redox signaling while exposing a collateral therapeutic vulnerability for a combination strategy. Our qHT-dS platform enables the accelerated discovery of Df-PPIs at single mutated residue resolution with the resultant dataset serving as a resource for the biomedical community to advance neoPPI candidates as targets for precision oncology approaches.

## RESULTS

### Cancer-associated gene expression library for focused PPI screening

Tumor evolution selects altered genes that provide fitness advantages to enable and sustain uncontrolled proliferation, progression, or invasion of cancer cells, which molds the landscape of cancer-associated proteins. To identify molecular re-wirings emanating from recurrently mutated residues encoded by genomic alterations, we sought to establish the connectivity of mutated residues and proteins with defined cancer-associated proteins for focused comparative examinations. For this purpose, we constructed a cancer-associated gene expression library, termed the OncoPPi v2 library (Fig. 1C & S2) (Li et al., 2017). Genes were selected based on their association with tumor pathogenesis in cancer patients, including gene amplifications, deletions, mutations, and rearrangements, across a wide range of tumor types (Fig. S2A; Table S1-2) (Cancer Genome Atlas Research et al., 2013; Forbes et al., 2017; Frampton et al., 2013; Lawrence et al., 2014). OncoPPi v2 library genes are significantly enriched in different oncogenic pathways, (e.g., *p*= 2.64e-94 for the “Pathway in cancer” gene set in KEGG database), cancer hallmarks (Fig. S2B), and defined oncogenes and tumor suppressors (Fig. S2C). The cancer-associated genes were cloned, sequence-verified, and used to create a collection of mammalian expression vectors with biosensor tags for 558 distinct human WT protein-coding open-reading frames (Table S1-2). The OncoPPi v2 library with significantly expanded genomic coverage was employed to search for interaction partners of driver mutations.

To select human cancer mutant alleles for Df-PPI discovery, we established a collection of oncogenic mutation expression vectors, termed the OncoMut library. For the systematic comparative study of driver mutations among oncogenes and tumor suppressor genes, we focused on recurrent mutations in two well-defined oncogenes and four frequently mutated tumor suppressor genes with a total of 24 alleles from diverse tumor lineages (Fig. 1D). These mutations were nominated based on their disease prevalence and their surface exposure potential for increased likelihood of creating neo-interaction epitopes (Kamburov et al., 2015). The OncoMut library contains mutations in serine/threonine-protein kinases AKT1 and BRAF and missense mutations in tumor suppressors, such as mutations in E3 ligase adaptors, Speckle type BTB/POZ protein (SPOP) and F-box and WD repeat domain containing 7 (FBXW7), and mutations in transcription regulators, mothers against decapentaplegic homolog 4 (SMAD4) and SWI/SNF related, matrix associated, actin dependent regulator of chromatin, subfamily A, member 4 (SMARCA4). The selected mutations that represent diverse protein classes in the OncoMut library (Fig. 1D), and together with the OncoPPi library (Fig. 1C), serve as the foundation for Df-PPI discovery.

### The qHT-dS platform for comparative WT and MUT PPI profiling

To systematically discover Df-PPIs in live cells, a quantitative high-throughput differential screening platform, qHT-dS, was established (Fig. 1B & S1). The qHT-dS platform is a highly efficient and quantitative approach enabled by using BRET^n^-based ultra-high throughput PPI screening (uHTS) technology in a 1536-well plate format (Mo and Fu, 2016; Mo et al., 2016) (Fig. S1A). Briefly, the expressed WT and MUT proteins, genetically tagged with Nanoluc-luciferase (NLuc) as a BRET^n^ donor, were systematically tested against each protein in the OncoPPi v2 library that are tagged with Venus as a BRET^n^ acceptor, in a parallel fashion (Fig. S1B). Due to the stringent proximity requirement (< 10 nm) to obtain a positive BRET^n^ signal, the identified PPIs generally reflect direct interactions in protein complexes (Mo and Fu, 2016; Mo et al., 2016). Upon substrate addition, three readouts were generated that capture: (i) luminescence from NLuc-fusions, (ii) fluorescence from Venus-fusion, and (iii) the ratiometric BRET signal from the Venus/NLuc emissions. The highly sensitive and multiplexed PPI signal detection of qHT-dS with simultaneous monitoring of test protein expression levels in a single add-and-read mode allows for large-scale screening of PPIs in live cells in a 1536-well plate format (Fig. S1C).

With the automated qHT-dS workflow (Fig. S1), we systematically tested 24 WT and MUT alleles for their binary interactions with each of the 558 cancer-associated proteins in the OncoPPi v2 library in HEK293T cells. A total of 13,392 interactions were examined with eight points of titration combinations in four replicates for each PPI’s BRET saturation curve (Table S3), generating a total of 1,474,560 data points. Such primary PPI signals were normalized to the luminescence signal for NLuc-tagged MUT or WT expression, and to the fluorescence signal for Venus-tagged OncoPPi genes expression. Ratiometric data for each PPI were calibrated with protein expression levels for each partner, enabling quantitative comparative assessment of each PPI from live cells, representing a differentiating factor from many other PPI HTS approaches. To ensure a rigorous statistical evaluation of the qHT-dS data, we developed a computational algorithm for automated Comparative Analysis of Rewired INterActions (CARINA). The CARINA algorithm operates directly with data in the 1536-well plate qHT-dS format and includes three major blocks of the analysis. First, saturation binding curves are calculated based on the qHT-dS data generated for the PPI and both corresponding empty donor/acceptor negative controls. Then, area under the curve (AUC) values are calculated to quantify the intensity of interaction signals (Fig. 2A). The fold-over-control (FOC) and *p*-value analysis of the AUC values (P_FOC_) derived from the BRET saturation curves are calculated to identify positive WT and MUT PPIs (Fig. 2B).

**Figure 2.**
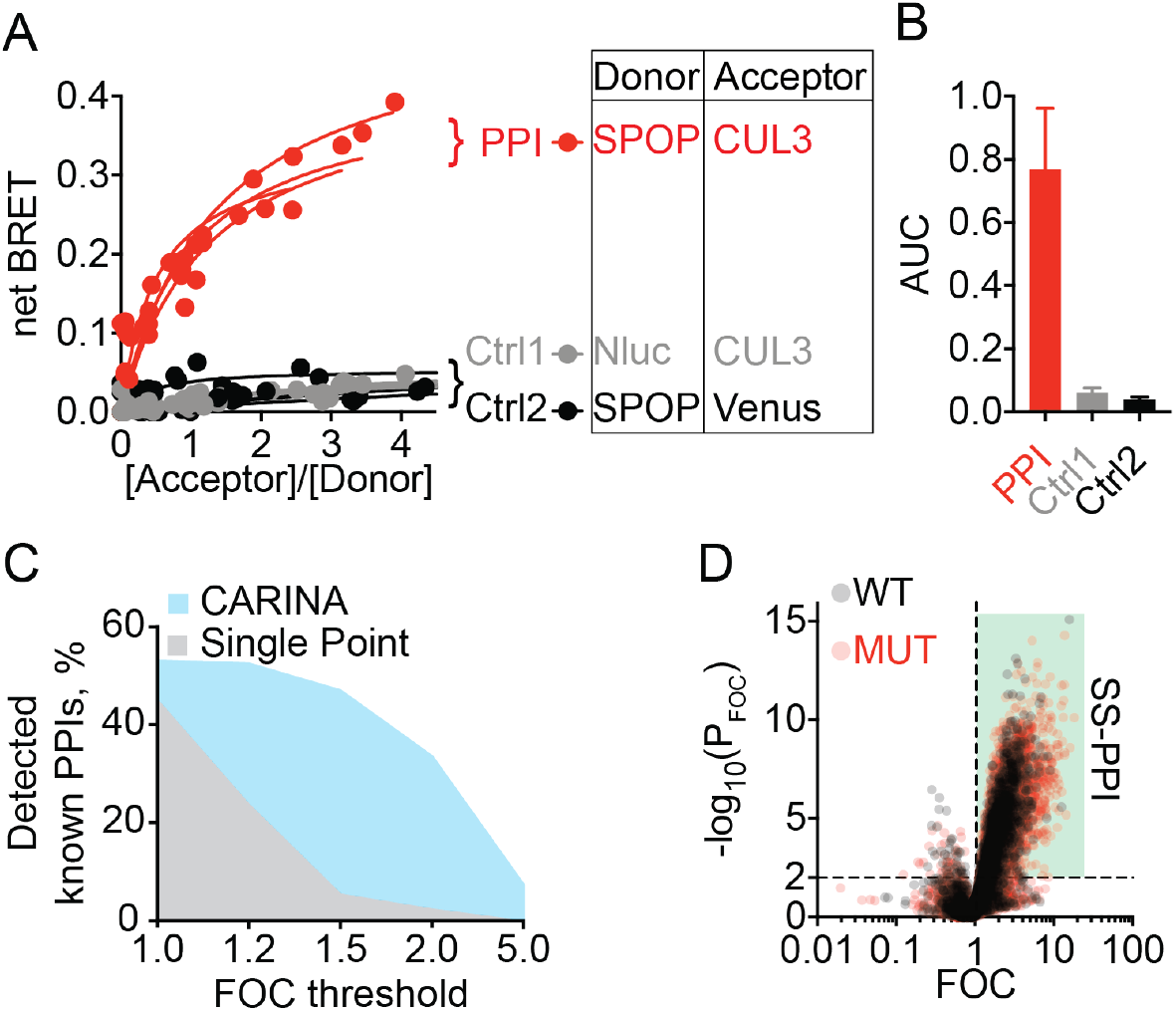
Evaluation of qHT-dS platform performance. **(A)** Representative BRET saturation curves from qHT-dS. Data from a known PPI, SPOP WT/CUL3, was used as an example and presented in four replicates. **(B)** AUC analysis of the BRET saturation curves using CARINA. **(C)** Positive discovery rate of the known WT PPIs using CARINA method comparing to the conventional single point analysis for the same dataset from qHT-dS. **(D)** Identification of statistically significant (SS) positive PPIs for both WT and MUT alleles.

Through CARINA analysis, a total of 40,176 BRET saturation curves were built for each PPI and its corresponding empty donor/acceptor controls. In contrast to conventional single point protein expression binary PPI mapping technologies (Li et al., 2017), qHT-dS incorporates variations in protein expression. As shown in Fig 2C, with a cut-off of FOC≥1.2 and P_FOC_≤0.01 CARINA identified more than 50% of known WT PPIs, while a lower FOC cut-off below 1.2 only slightly increased the number of known PPI detected in the screening. Therefore, FOC≥1.2 and P_FOC_≤0.01 were used as the primary threshold to define statistically significant (SS) PPIs. With these parameters, a total of 6,190 primary positive PPIs were identified (Fig. 2D, Table S4). These positive PPIs serve as a candidate pool for further evaluation to identify differential PPIs.

### Systematic analysis to prioritize differential PPIs

To identify mutant-driven Df-PPIs, we performed the comparative analysis according to the FOC values of WT and MUT PPI profiles. To differentiate gain of interactions (Go-PPI) from loss of interactions (Lo-PPI), the difference between WT and MUT PPI curves were calculated as the ratio of the averaged AUC obtained for the MUT-PPI over the WT-PPI and quantified as differential scores (DS) with the corresponding DS T-test p-values (P_DS_) (Fig. 3A, Table S4). Based on DS and P_DS_ cut-offs, 889 differential Go-PPI (DS>1 and P_DS_≤0.001) and 161 differential Lo-PPIs (DS<1, and P_DS_≤0.001) were identified. To prioritize PPIs for confirmatory studies, the adjusted p-value (Q_DS_) cut-off of Q_DS_≤0.01 and high stringent (HS) cut-offs of DS≥1.5 or DS≤1/1.5, indicating at least 50% difference between WT and MUT PPI signals, were applied to compile the HS-Go-PPI and HS-Lo-PPI groups of 409 and 55 PPIs, respectively (Fig. 3A-B, Table S4). These 464 HS-differential PPIs were utilized to assess the specificity and nature of allele-dependent interactions.

**Figure 3.**
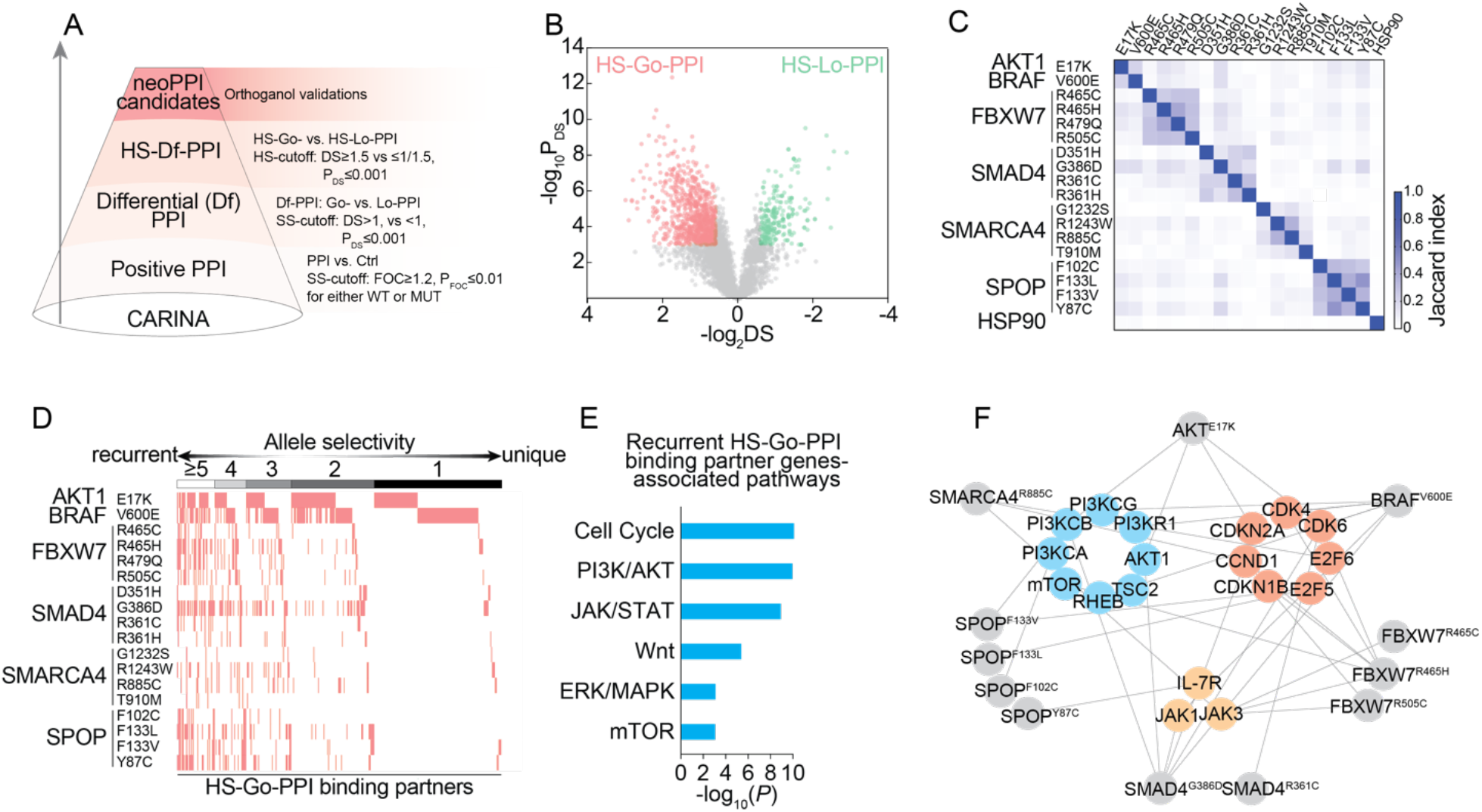
Systematic analysis to prioritize differential PPI to advance neoPPI candidates. **(A)** Funnel chart of CARINA for differential PPI identification and neoPPI candidate prioritization with cut-off criteria **(B)** Volcano plot of the differential score versus p values for each pairwise WT/MUT PPI shows the identification and prioritization of HS-Go- and HS-Lo-PPI. **(C)** Similarity analysis of HS-Go-PPI binding partners between alleles. **(D)** Clustering analysis reveals recurrent and unique HS-Go-PPI binding partners. **(E)** The commonly re-wired pathways that are significantly enriched from the pathway analysis of the recurrent HS-Go-PPI binding partners. **(F)** Go-PPI network of oncogenic pathways that are commonly re-wired by various driver mutations of diverse genes. Each neoPPI is double positive from both qHT-dS and GST pull-down validation.

Different genes, or even different alleles from the same gene, showed remarkable divergence in their interaction partners from an analysis of their HS-Go-PPI profiles (Fig. 3C). For example, less than 10% of overlapping Go-PPI partners were found for mutations in two E3 ligase substrate recognition proteins, SPOP and FBXW7. The fraction of shared partners for G386D vs. D351H SMAD4 alleles, and F102C vs. F133V SPOP alleles was 13% and 30%, respectively, exhibiting mutation-specific interactions. Surprisingly, mutations at the same site showed differential interactions in a residue-dependent manner. As an example, 62% of SPOP F133L partners were different from those of F133V. Such mutant-allele specific interactions were unlikely due to association with the general chaperone function of heat shock protein 90 (HSP90) even though a number of disease mutants show increased interaction with HSP90 (Oughtred et al., 2018; Sahni et al., 2015). Indeed, only minimal overlap (1.4±0.9%) between HS-Go-PPI and HSP90 known PPI partners was noted (Fig. 3C). These results highlight the specificity of mutant allele-mediated Go-PPIs and potential allele-dependent tumor heterogeneity at the protein connectivity level.

To gain insights into allele-driven oncogenic programs, allele selectivity for each partner was examined from a clustering analysis (Fig. 3D). Allele selectivity revealed that 35% HS-Go-PPI partners interact with 3 or more alleles, implicating shared oncogenic pathways by different mutations. The gene ontology and pathway analysis revealed that the recurrent HS-Go-PPI partners function in core growth regulatory pathways, including cell cycle, PI3K/AKT/mTOR, and JAK/STAT (Fig. 3E, Table S5). To verify such mutated residue-mediated interactions with core mediators of oncogenic pathways, we selected three representative pathways and carried out orthogonal affinity-based GST pull-down assay. As illustrated in Fig. 3F, a commonly re-wired pathway network was compiled based on the validated Df-PPIs, showing the engagement of various mutated residues on common pathways, such as cell cycle, PI3K/AKT/mTOR and JAK/STAT pathways, at different signaling protein levels (Fig. S3). Additionally, the majority of HS-Go-PPI partners (65%) interact with one or two alleles, suggesting that these unique partners and corresponding pathways could be specifically altered in cancers harboring certain mutations, or represent new mechanisms that engage core oncogenic programs (Fig. 3D).

### neoPPI candidates induced by mutations of tumor suppressors and oncogenes

To gather further experimental evidence to advance the characterization of mutated allele-directed differential PPIs, we sampled HS-Df-PPIs from the qHT-dS BRET dataset to compare their interactions in an affinity chromatography-based GST pull-down assay that involved multiple washing steps. Comparative affinity pull-down studies were carried out to evaluate the interaction of mutant allele-mediated PPIs relative to that of WT counterparts. As shown in Fig. 4 and Fig. S3, 265 out of 325 PPIs were found to show differential binding signals, representing 82% of HS-Df-PPIs with supporting evidence from both BRET and GST pull-down assays. These double positive Df-PPIs include mutant-mediated Go-PPIs for both oncogenes and tumor suppressors, which were used as mutation-enabled neoPPI candidates for functional evaluation.

**Figure 4.**
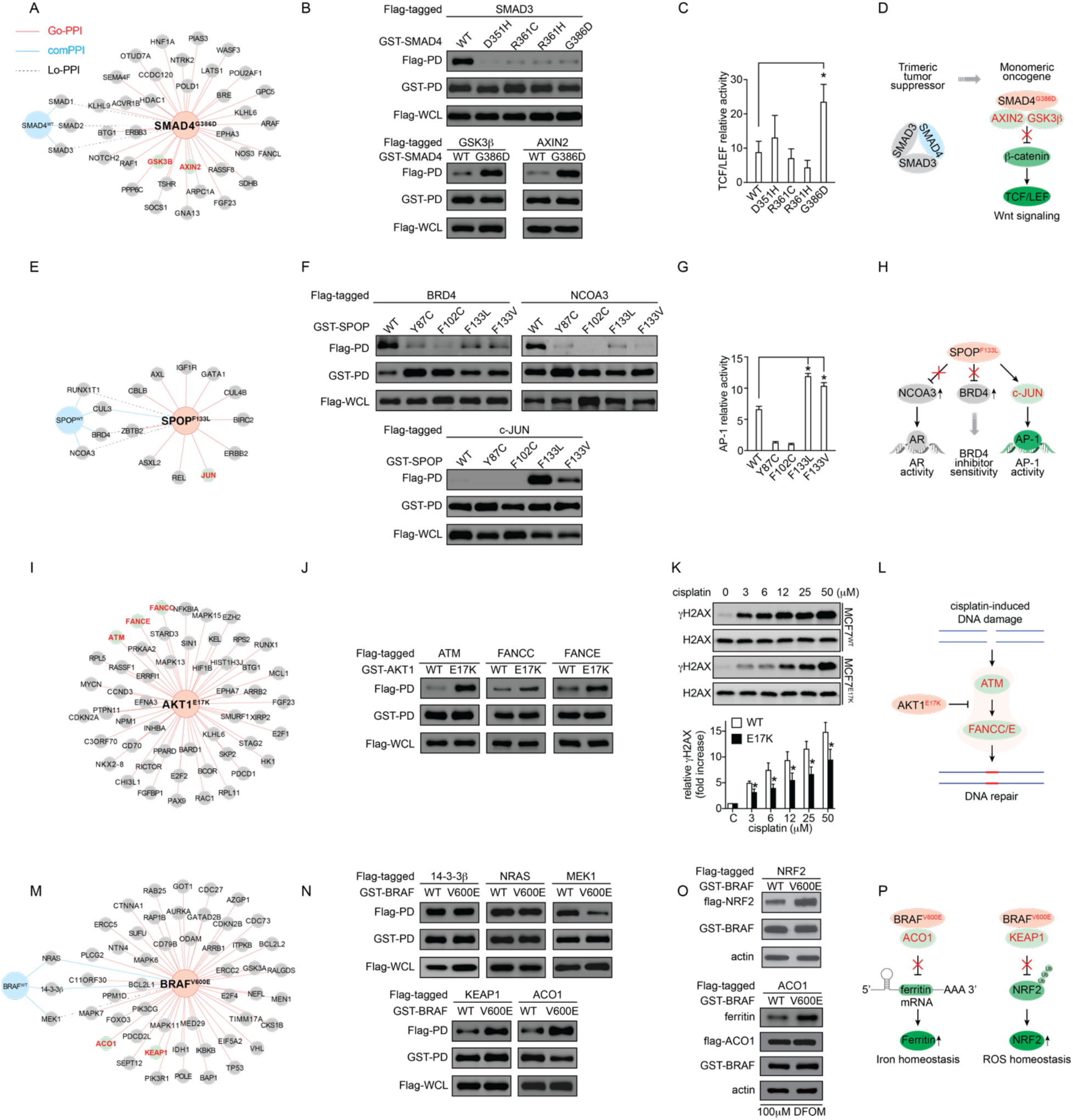
neoPPI candidates induced by mutations of tumor suppressors and oncogenes. See also Figure S3-4. **(A, E, I, and M)** Hub and spoke diagram of four major neoPPI hubs revolved around **(A)** SMAD4^G386D^, **(E)** SPOP^F133L^, **(I)** AKT1^E17K^ and **(M)** BRAF^V600E^. **(B, F, J and N)** GST pull-down validation results of the selected Go-PPI, comPPI and/or Lo-PPI in HEK293T cell overexpressing the construct as indicated. Confirmation of **(B)** SMAD4^G386D^’s Lo-PPI with SMAD3, and neoPPIsneoPPIs with GSK3β and AXIN2, **(F)** SPOP^F133L^’s Lo-PPI with BRD4 and NCOA3, and neoPPI with c-JUN **(J)** AKT1^E17K^’s neoPPIs with ATM, FANCC and FANCE, **(N)** BRAF^V600E^’s comPPIs with 14-3-3β 3β and NRAS, Lo-PPI with MEK1, and neoPPIs with KEAP1 and ACO1. **(C, G, K and O)** Functional examination of the selected neoPPIs. **(C)** TCF/LEF transcriptional reporter activity in HEK293T cells transfected with GST-SMAD4^G386D^ or other mutants comparing to the WT controls. The data are presented as mean ± SD from three independent experiments. *p≤0.05. **(G)** AP-1 transcriptional reporter activity in HEK293T cells transfected with GST-SPOP^F133L^ or other mutants comparing to the WT controls. The data are presented as mean ± SD from three independent experiments. *p≤0.05. **(K)** Western blots showing the γH2AX and total H2AX levels in MCF7 WT (PI3KCA^WT^/AKT1^WT^) and MCF7 E17K (PI3KCA^WT^/AKT1^E17K^) upon treatment with cisplatin at concentration as indicated. The relative γH2AX was calculated as the fold induction of γH2AX/H2AX comparing to no cisplatin control. The data are presented as mean ± SD from three independent experiments. *p≤0.05. **(O)** Western blots showing the NRF2 protein levels in HEK293T cells co-transfected with GST-BRAF^V600E^ and flag-tagged NRF2 comparing to the WT control (*upper*), and the ferritin protein levels in HEK293T cells co-expressing GST-BRAF^V600E^ and flag-ACO1 in the presence of 100 μM deferoxamine mesylate (DFOM) (*lower*). **(D, H, L and P)** Hypothetical model for further interrogation of neoPPI-induced pathway reprograming. **(D)** Model of SMAD4^G386D^/AXIN2 and GSK3β neoPPIs in activating Wnt/β-catenin signaling pathways. **(H)** Model of SPOP^F133L^/c-JUN neoPPI in regulating AP-1 transcriptional activities. **(L)** Model of AKT1^E17K^/ATM and FANCC/E neoPPIs in regulating cisplatin-induced DNA damage response pathway. **(P)** Model of BRAF^V600E^/ACO1 and KEAP1 neoPPIs in reprogramming iron and ROS homeostasis pathways.

### Mutation effect of tumor suppressor genes

Tumor suppressor genes were generally believed to enable tumorigenesis through loss-of-function mutations, which impair their normal tumor-suppressive functions(Cheng et al., 2021). Unexpectedly, the Df-PPI data showed that tumor suppressor mutations exhibited both Lo-PPIs and Go-PPIs with a different set of cancer-associated proteins. Two representative hubs (Fig. 4), centered around SMAD4^G386D^ and SPOP^F133L^, were identified with their allele selective partners supported by the physical interaction validation.

SMAD4 is a core intracellular effector of the canonical Transforming Growth Factor-β (TGF-β) signaling pathway (Shi and Massague, 2003; Wrana, 2009). Upon TGF-β activation, SMAD4 forms a trimeric complex with receptor-activated SMADs (SMAD1/2/3) that translocate into the nucleus for transcriptional regulation. SMAD4 mutations in the MH2 domain that disrupt the trimeric complex have been identified in various tumor types (Chacko et al., 2004; Miyaki and Kuroki, 2003). Indeed, our qHT-dS platform recapitulated the known Lo-PPI of SMAD4 mutants, such as D351H, R361C/H and G386D, with SMAD3 (Fig. 4A-D), supporting their known monomeric state (Chacko et al., 2004). We also uncovered new Lo-PPIs binding partners, including LZTR1 and GATA2, that are positive for SMAD4 WT but weakened upon mutations (Fig. S3A). These Lo-PPIs may serve as potential molecular targets to develop hypomorph mutation-directed PPI inducers (Tang et al., 2020).

Interestingly, SMAD4^G386D^ exhibited enhanced interactions with 34 binding partners (Fig. 4A, Fig. S3A), 25 of which had added evidence of co-localization in intracellular compartments with SMAD4 (Table S6). These results suggest that SMAD4^G386D^ may have potential non-canonical properties in regulating the corresponding pathways in addition to its known loss-of-interactions. As an example, SMAD4^G386D^ showed significantly (p-value < 0.05) elevated interaction with AXIN2 and GSK3β (Fig. 4B), two core components in the Wnt signaling pathway, leading to the hypothesis that SMAD4^G386D^ may gain an ability to impact AXIN2 and GSK3β−regulated β-catenin/TCF transcriptional activity. Indeed, cells overexpressing SMAD4^G386D^ showed significantly higher TCF/LEF transcriptional luciferase reporter signals than that induced by SMAD4 WT or other mutants (p-value < 0.05, Fig. 4C). These results implicated the G386D induced transition from the SMAD4/SMAD3 trimeric tumor suppressor complex to a new protein complex that enables SMAD4 crosstalk with the Wnt signaling pathway through SMAD4 mutant-mediated neoPPIs (Fig. 4D).

SPOP is an adaptor protein in CUL3-SPOP E3 ubiquitin ligase complexes that mediates substrate binding for ubiquitination and degradation (Zhuang et al., 2009). Genomic studies have revealed missense mutations in SPOP’s substrate-recognition MATH domain that recur in prostate cancers (Barbieri et al., 2012). Several SPOP mutations in the MATH domain have been defined as loss-of-function that lead to loss of oncogenic substrate recognition, reduce substrate ubiquitination, and promote aberrant elevation of substrate protein levels. Collectively, this suggests that SPOP WT has tumor suppressive functions that are lost through recurrent missense mutations (Dai et al., 2017; Janouskova et al., 2017; Theurillat et al., 2014; Zhang et al., 2017). In support of this notion, from the qHT-dS (Fig. 4E-H), we identified several previously characterized SPOP WT interaction partners, including and BRD4 and NCOA3 (SRC-3), as Lo-PPIs for all four tested SPOP mutations, supporting their loss-of-function role (Fig. 4F). In addition, we also identified and validated new SPOP WT binding partners, such as RUNX1T1 and ETV1, that exhibit loss of interaction with mutant SPOP variants (Fig. S3B). These results further support the current loss-of-function annotation of SPOP mutations and expanded the SPOP WT-PPI spectrum.

Unexpectedly, the qHT-dS screening uncovered a panel of Go-PPIs as neoPPI candidates that were associated with mutated SPOP, such as SPOP^F133L^. The F133L mutation not only disrupted SPOP interaction with known SPOP substrates, but also gained the ability for enhanced interaction with thirteen new partners (Fig. 4E, Fig. S3B). As an example, SPOP^F133L^ showed increased interaction with c-JUN (Fig. 4F), a protein in combination with c-FOS forming the AP-1 early response transcription factor, leading to the hypothesis that SPOP^F133L^ may modulate AP-1 transcriptional activity. To test this notion, we examined AP-1 transcriptional activity and observed that HEK293T cells expressing SPOP^F133L^ have significantly increased AP-1 reporter activity (p-value < 0.05) compared to that of WT SPOP (Fig. 4G). These results not only support previously reported roles of SPOP mutations in lost tumor suppressor activity by releasing bound oncoproteins, but also suggest potential JNK-c-JUN pathway re-wiring induced by the previously unrecognized neoPPI, SPOP^F133L^/c-JUN (Fig. 4H).

Such gain-of-interaction features were not restricted to SMAD4 and SPOP; we also detected neoPPI candidates for other tumor suppressor mutations, such as FBXW7 and SMARCA4. For example, we have identified and confirmed several FBXW7 and SMARCA4 mutation-mediated PPIs with binding partners, such as CCND1, and JAK3, with the GST pull-down assay, suggesting their potential connectivity in regulating the cell cycle, PI3K/mTOR and JAK/STAT pathways (Fig. 3 and Fig. S3C). In addition, we detected FBXW7 and SMARCA4 mutation-selective neoPPIs, such as FBXW7^R465H^/C8ORF4, FBXW7^R465H^/MKK3B, and SMARCA4^R885C^/B2M (Fig. S3C), implying a potential role of the FBXW7^R465H^ in C8ORF4 regulated Wnt and NOTCH pathway (Jung et al., 2006; Zhu et al., 2015), or in MKK3B-centered MAPK signaling (Wagner and Nebreda, 2009); and the SMARCA4^R885C^ in B2M dependent antigen presentation machinery (Restifo et al., 1996).

Altogether, our data provide a proof-of-concept that these previously annotated “loss-of-function” tumor suppressor mutations may have unexpected neomorphic activity in engaging new binding partners. These tumor suppressor variant-driven neoPPIs illustrate another dimension of complexity underpinning the role of tumor suppressor in enabling tumorigenesis and cancer progression. Both neoPPIs and Lo-PPIs identified and confirmed from our study may be targeted for therapeutic intervention through the discovery of neoPPI inhibitors or hypomorph-PPI inducers (Tang et al., 2020).

### Mutation effect of oncogenes

Dysregulated kinases are a major family of oncogenes and attractive druggable targets. Kinase inhibitors have been developed and shifted the paradigm of oncology practice towards precision oncology. However, drug resistance significantly limits the impact of oncogenic kinase inhibitors. Such clinical challenges demand better understanding of the molecular underpinning of tumorigenesis driven by “gain-of-function” kinase mutations. Our qHT-dS data showed that driver mutations enhanced the interaction of respective kinases, such as AKT1^E17K^ and BRAF^V600E^, with new binding partners, which may re-wire the oncogenic pathways and reshape therapeutic responses (Fig. 4).

AKT1 is a serine-threonine kinase that is frequently activated in a wide range of cancers (Porta et al., 2014). E17K in the Pleckstrin homology domain of AKT1 is the most frequent hot-spot gain-of-function mutation, leading to its recruitment to the plasma membrane and activation of downstream signaling (Carpten et al., 2007; Hyman et al., 2017; Jung et al., 2016). From our PPI screening, AKT1^E17K^ appeared to form a major hub with 51 allele selective binding partners (Fig. 4I, Fig. S3D). Among these 51 binding partners, 47 co-localize in AKT1’s known cellular compartments, 11 were known AKT1 substrates or WT binding partners, and 14 were novel AKT1^E17K^ binding partners containing the defined AKT1 phosphorylation consensus motif RxRxxS/T in support of their association with AKT1 (Table S7). To examine the functional implication of AKT1^E17K^-enhanced PPIs, we selected the E17K-enhanced interaction of AKT1 with ATM for further analysis. ATM is involved in DNA damage response pathways (Fig. 4J) (Ceccaldi et al., 2016; Shiloh and Ziv, 2013). This AKT1^E17K^/ATM neo-interaction may alter response to DNA damage and repair. To test this notion, we examined Histone H2AX phosphorylation (γH2AX), a marker for DNA damage (Sharma et al., 2012), in a pair of isogenic AKT1 WT and E17K MCF7 cell lines (Beaver et al., 2013) after treatment with cisplatin, a chemotherapeutic agent known to induce DNA damage (Dasari and Tchounwou, 2014). Upon cisplatin treatment, we detected a dose-dependent increase of γH2AX in AKT1^E17K^ cells, but to a significantly less extent in the isogenic AKT1 WT MCF7 cells (Fig. 4K). Besides ATM, AKT1^E17K^ partners also include FANCC and FANCE, two additional factors in the DNA damage response pathway (Fig. 4J). These results provide a potential connectivity of AKT1^E17K^ with ATM and FANCC/E and their functional roles in regulating the DNA damage response (Fig. 4L). This neoPPI-informed model is consistent with the previously recognized association between AKT1 activation and DNA damage and repair (Liu et al., 2014; Oeck et al., 2017), and provides potential mechanistic underpinnings for this association.

BRAF is a core component of the RAS/RAF/MEK/ERK signaling cascade, a pathway that is frequently dysregulated in multiple tumor types (Ascierto et al., 2012; Cancer Genome Atlas Research, 2014; Cantwell-Dorris et al., 2011; Karoulia et al., 2017). The hot-spot mutation, V600E, in the activation segment of the BRAF kinase domain results in kinase hyperactivation and dysregulation of MEK/ERK signaling. From qHT-dS and GST pull-down validation (Fig. 4M-N, Fig. S3E), we confirmed common PPIs with RAS and 14-3-3β, and Lo-PPI with MEK1 for BRAF^V600E^ compared to the WT counterpart, supporting the reported model of BRAF^V600E^ in regulating the canonical RAS/RAF/MEK pathway signaling (Haling et al., 2014). We also identified and confirmed 48 V600E-enhanced neoPPI candidates, significantly expanding the BRAF^V600E^-mutation allele-mediated oncogenic pathways beyond the RAS/RAF/MEK1-centered signaling (Fig. 4M-N, Fig. S3E).

Next, we assessed the potential biological relevance of the BRAF^V600E^-induced neo-interactions with database mining and annotation. First, bioinformatics analysis revealed that alterations in 45 out of 48 neoPPI binding partners showed mutual exclusivity with BRAF^V600E^ in thyroid, melanoma, or colon cancers, and 39 binding partners co-localize with BRAF, supporting their functional associations (Fig. S4, Table S8). Second, we examined the potential correlation of these PPIs with tumor proliferation and survival. We reasoned that if a binding partner is functionally important for the proliferation of BRAF^V600E^-harboring cells, the perturbation of the binding partner gene expression would generate greater effects on survival in V600E than that in WT cells. From the analysis of gene knockdown/knockout screen dataset from The Cancer Dependency Map (DepMap), V600E harboring cells showed significantly lower CERES/DEMETER scores than that of WT cells (p-value ≤ 0.05) upon knocking-down/out of selected V600E-neoPPI partners, which include AZGP1, BAP1, EMSY, KEAP1, MAPK11, NTN4, PDCD2L, PIK3CG, PPM1D and TIMM17A (Table S8). These datasuggest the potential functional significance of these neoPPIs in maintaining V600E-cell proliferation and survival. Lastly, we considered the potential correlation of these neoPPIs with therapeutic response of V600E-harboring tumors. It is possible that expression levels of binding partners might be correlated with sensitivities of V600E-cells to gene perturbation and drug treatment. To test this notion, we correlated the mRNA or protein expression of binding partners with their associated sensitivity scores in response to drug treatment. From the analysis of therapeutic response profiles of the CTRP dataset, the drug sensitivity of V600E harboring cell lines to vemurafenib, a clinically approved BRAF^V600E^ inhibitor, showed statistically significant correlation (Pearson correlation coefficient |R|≥0.3 and p-value ≤ 0.05) with mRNA or protein expression levels of 14 genes, such as aconitase 1 (ACO1) and BCL2L1 (Table S8). These data add evidence in support of the functional importance of BRAF^V600E^-neoPPIs for tumor proliferation, survival or therapeutic response.

To further gather evidence to probe the functional importance of BRAF^V600E^ neoPPI candidates, we selected the interaction of BRAF^V600E^ with ACO1 and the Kelch-like ECH-associated protein 1 (KEAP1) (Fig. 4N) for experimental evaluation. ACO1, also known as Iron-Responsive Element-Binding Protein 1 (IRP1), is a bifunctional protein that regulates iron homeostasis through regulating ferritin levels (Beinert et al., 1996). KEAP1 is an adaptor protein in E3 ligase complexes that mediate ubiquitination and degradation of protein substrates, such as NF-E2-related factor 2 (NRF2), a master regulator of reactive oxygen species (ROS) homeostasis (Jaramillo and Zhang, 2013). We observed that cells expressing BRAF^V600E^ showed significantly increased ferritin and NRF2 protein levels compared to that of BRAF WT (Fig. 4O). These results suggest that BRAF^V600E^ may reprogram the iron and ROS homeostasis through the neoPPIs with ACO1 and KEAP1, respectively (Fig. 4P).

### Neo-interaction of BRAF^V600E^ with KEAP1 and its collateral vulnerability

To illustrate the utility of the discovered neoPPI candidates to inform oncogenic mechanisms for potential intervention strategies, we selected and examined the neo-interaction of BRAF^V600E^ with KEAP1. The mutated BRAF^V600E^ created a neo-epitope with enhanced affinity to KEAP1, which may affect ROS response through modulating the NRF2-mediated transcriptional and metabolic machinery (Jaramillo and Zhang, 2013; Sporn and Liby, 2012). To test this hypothesis, we extensively assessed this neoPPI with a panel of biochemical and cellular assays.

Consistent with the qHT-dS readouts (Fig. 5A), BRAF^V600E^ showed significantly higher interaction signals than the WT with KEAP1 in an orthogonal GST pull-down assay (Fig. 4N). This neo-interaction was further demonstrated in live cells showing cytoplasmic localization with the Venus-PCA assay (Fig. 5B), and under endogenous cellular conditions with co-immunoprecipitation studies in a pair of isogenic cell lines (Fig. 5C) and a panel of patient derived BRAF^V600E^ melanoma cell lines (Fig. S5), supporting its presence under physiological conditions. Truncation studies suggested the involvement of the kinase domain of BRAF^V600E^ and the KELCH domain of KEAP1 in their association (Fig. 5D). This neo-interaction appears to be direct and reversible. With the purified proteins in a Biolayer Interferometry (BLI) analysis, the kinase domain of BRAF^V600E^, but not that of WT, exhibited direct binding BLI signals to the KEAP1 KELCH domain (Fig. 5E). Treatment of BRAF^V600E^ carrying cells with vemurafenib, an FDA approved BRAF^V600E^ kinase inhibitor, attenuated the BRAF^V600E^-KEAP1 interaction in a dose-dependent manner (Fig. 5F). Together, these data strongly support a physiologically relevant neoPPI, BRAF^V600E^-KEAP1.

**Figure 5.**
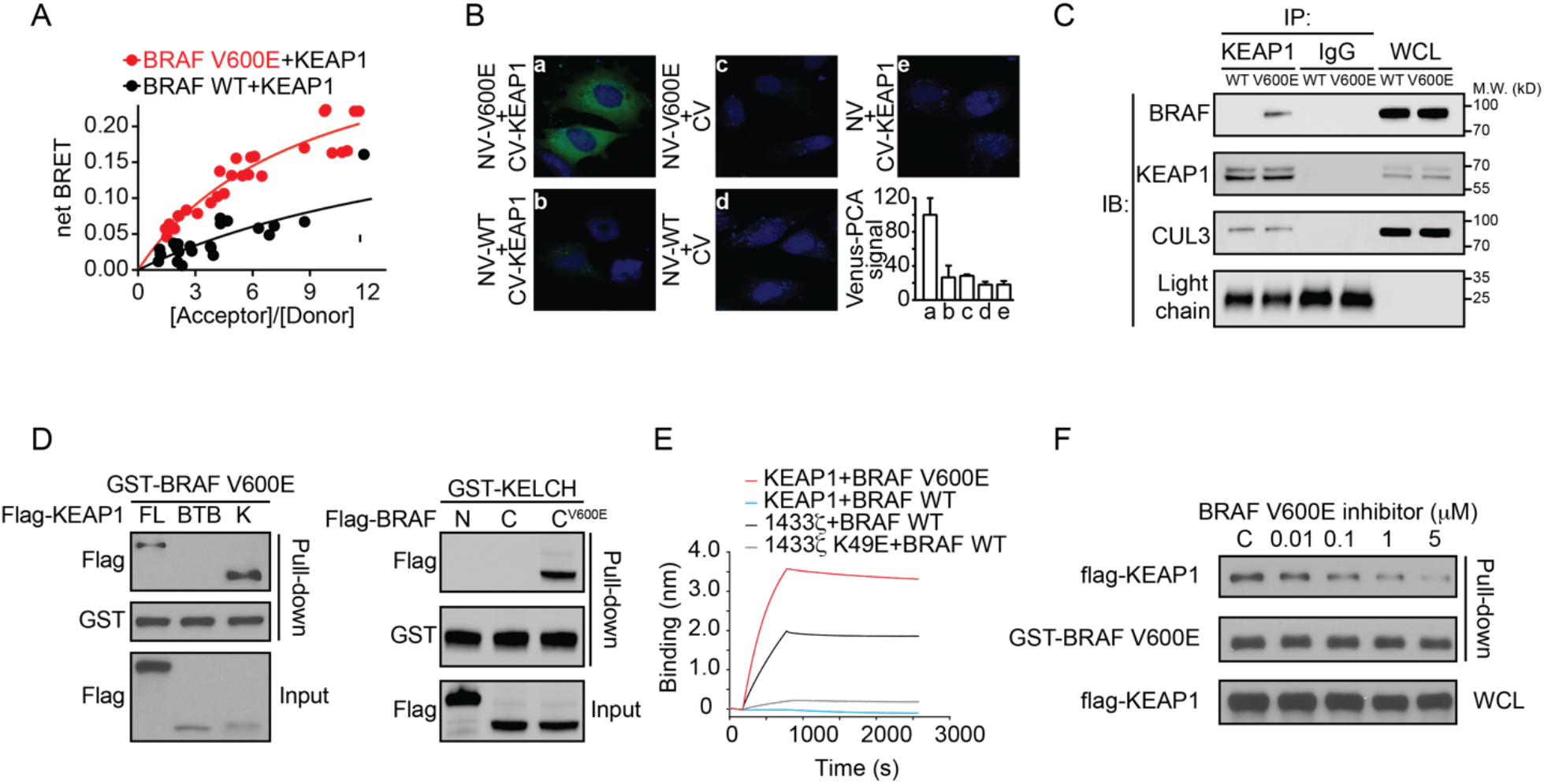
Validation of BRAF^V600E^ interaction with KEAP1. See also Figure S5. **(A)** BRET^n^ saturation curve of Venus-tagged KEAP1 interaction with NLuc-tagged BRAF WT versus V600E from qHT-dS. The data is presented by combining four replicates from the primary qHT-dS. **(B)** Venus-PCA shows the cytoplasm localization of BRAF^V600E^/KEAP1 neoPPI using CHL-1 melanoma cell line transfected with N-Venus-tagged BRAF WT or V600E and C-Venus-tagged KEAP1. Green: reconstituted Venus signal. Blue: nuclear stained with Hoechest. Venus-PCA signal was presented as the normalized fluorescence intensity. **(C)** Endogenous interaction of BRAF^V600E^ with KEAP1. The BRAF^V600E^/KEAP1 complex was co-immunoprecipitated with KEAP1 antibody from a pair of isogenic colon cancer cells, RKO and RKO (+/-/-), with anti-IgG as control. **(D)** GST-affinity pull-down assay with BRAF and KEAP1 fragments suggests the interaction domains including KEAP1 KELCH domain (left panel) and BRAF^V600E^ kinase domain (right panel). **(E)** Bio-layer interferometry assay validation of the direct interaction between KEAP1 KELCH domain and BRAF^V600E^ kinase domain using human recombinant proteins. Interaction between 14-3-3ζ and BRAF WT kinase domain was used as positive control, and 14-3-3ζ K49E and BRAF WT were used as negative control. **(F)** Effect of BRAF^V600E^ kinase inhibition by vemurafenib on BRAF^V600E^/KEAP1 neoPPI. HEK293T cells transfected with GST-BRAF^V600E^ and flag-KEAP1 were treated with vemurafenib at indicated concentrations for six hours.

These data suggested an undescribed function of the mutated BRAF as a regulator of the KEAP1-mediated redox pathway by directly impinging on the KEAP1-NRF2-ARE signaling axis. Indeed, overexpression of BRAF^V600E^, but not WT, significantly stabilized NRF2 protein (Fig. 6A-B), increased its transcriptional ARE reporter activity (Fig. 6C), and increased protein levels of NRF2 and its target gene, NQO1 (NAD(P)H Quinone Dehydrogenase 1)(Fig. 6D), but not NRF2 mRNA levels (Fig. 6E). NRF2 protein and its downstream NQO1 protein levels were significantly elevated in a panel of BRAF^V600E^ human melanoma cell lines, as compared to that of the WT cell lines (Fig. 6F-G). These data were further supported by the large-scale CCLE dataset, showing a positive correlation of upregulated NQO1 mRNA level with the V600E status of BRAF (Fig. 6H, Table S9). Further, treatment of BRAF^V600E^ melanoma cells with vemurafenib that decreased KEAP1 binding led to downregulated NRF2 activity and NQO1 protein levels (Fig. 5F and Fig. 6I). The MEK1 inhibitor had no effect on the KEAP1-NRF2 interaction (Fig. 6J). To determine how BRAF^V600E^ regulates the KEAP1-NRF2 signaling, we carried out a PPI competition experiment. Increased expression of BRAF^V600E^ was correlated with the reduced amount of NRF2 in the KEAP1 complex, suggesting the displacement of NRF2 from the KEAP1 complex by BRAF^V600E^ (Fig. 6K). These results suggest a BRAF^V600E^-induced KEAP1 sequestration model in which BRAF^V600E^ activates NRF2 through a competitive binding with KEAP1, in addition to the previously reported transcriptional regulation mechanism (DeNicola et al., 2011).

**Figure 6.**
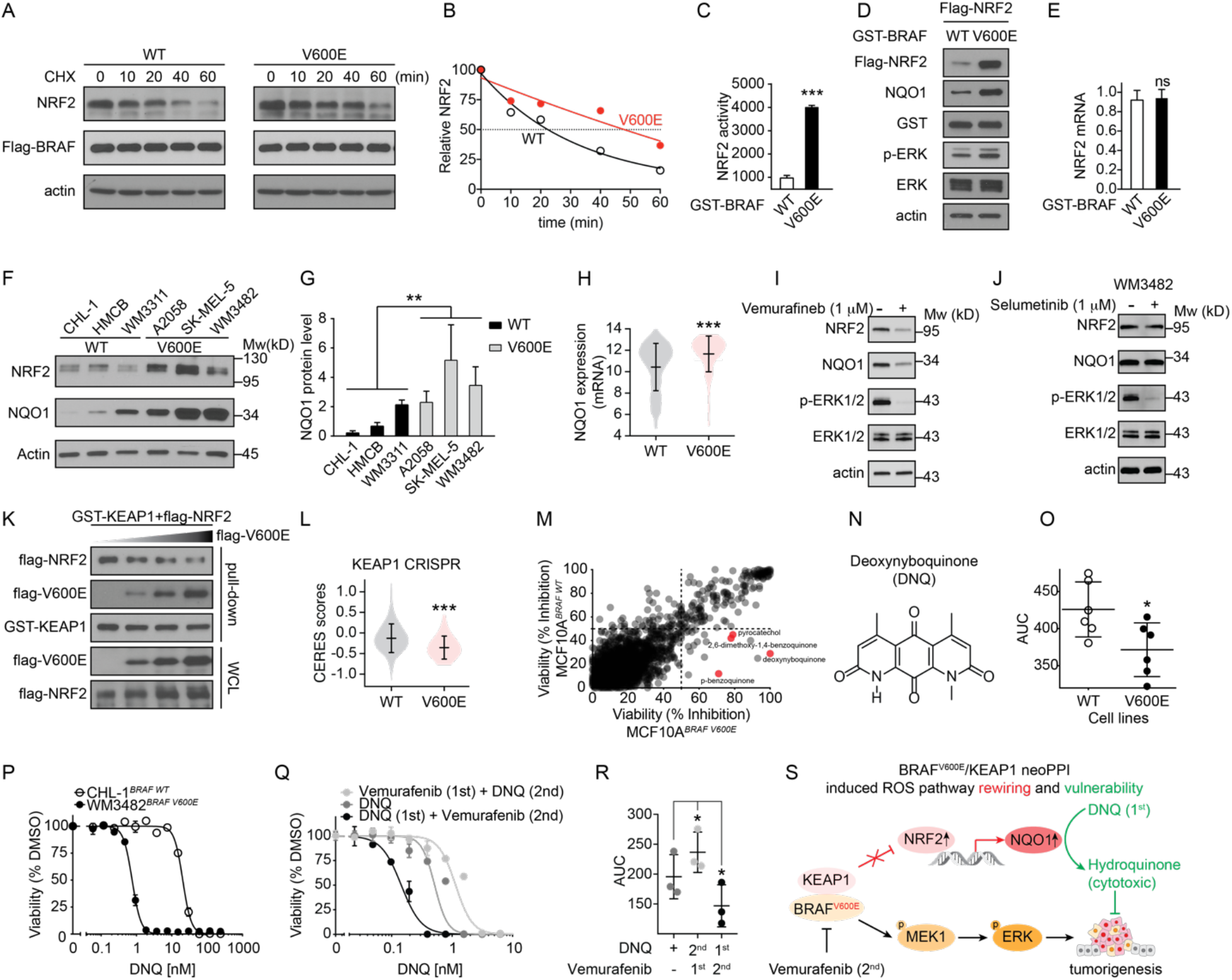
Neo-interaction of BRAF^V600E^ with KEAP1 and its collateral vulnerability. **(A)** BRAF^V600E^ stabilizes endogenous NRF2. Immunoblot showing NRF2 and actin levels at different time points after inhibition of protein synthesis with cycloheximide in HEK293T cells overexpressing BRAF WT or V600E. **(B)** Graph of NRF2 protein levels at indicated time points based on densitometric analysis of results in (A). **(C)** BRAF^V600E^ activates NRF2 transcriptional activity. HEK293T cells were co-transfected with the NRF2-ARE luciferase reporter and either WT or V600E BRAF. Relative luciferase activity was measured, normalized to internal Renilla luciferase control. Representative results of three independent experiments are shown. The error bars show the mean ± SD of three replicates. ***p<0.001. **(D)** BRAF^V600E^ increases NRF2 and its target gene NQO1 protein levels in HEK293T cells transfected with GST-BRAF V600E versus WT. **(E)** Effect of BRAF^V600E^ on NRF2 mRNA levels in HEK293T cells transfected with flag-NRF2 and GST-BRAT WT or V600E plasmids. ^ns^p>0.05. **(F)** Correlation of NRF2 and its target gene NQO1 protein levels with BRAF genetic status in six melanoma cell lines with WT or V600E BRAF. **(G)** Bar graph of NQO1 protein levels in indicated cell lines based on densitometric analysis of results in (F). **p<0.01. **(H)** Violin plot of the correlation between NQO1 mRNA levels and BRAF genetic status in 967 cell lines from the Cancer Cell Line Encyclopedia genomic dataset. The lines indicate mean and SD. ***p<0.001. **(I-J)** Effect of BRAF^V600E^ kinase inhibitor, vemurafenib **(I)**, or MEK1 kinase inhibitor, selumetinib **(J)**, on NRF2 and NQO1 protein levels in WM3482 melanoma cell line with BRAF V600E mutation. **(K)** Competitive binding between BRAF^V600E^ and NRF2 to KEAP1. GST-affinity pull down of GST-KEAP1 complex from lysate of HEK293T cells transfected with equal amounts of GST-KEAP1 and flag-NRF2, and with increasing amounts of flag-BRAF^V600E^ plasmid. **(L)** Violin plot showing the CERES scores for CRISPR knockout of KEAP1 in 342 cancer cell lines from CCLE dataset. The lines indicate mean and SD. ***p<0.001. **(M)** Parallel HTS of chemical genomic compound library in a pair of isogenic MCF10A cell lines. The identification of quinone and its derivatives with selective growth inhibition of cell line with BRAF^V600E^ are highlighted in red. Data were presented as percentage of inhibition in parental MCF10A cells with BRAF WT versus its V600E knock-in counterpart for 3200 bioactive compounds. **(N)** Chemical structure of one of the quinones, deoxynyboquinone (DNQ). **(O)** AUC analysis of DNQ-induced dose-dependent growth inhibition of twelve cell lines with BRAF WT or V600E. WT: CHL-1, HMCB, MCF10A, MeWO, WM3311 and RKO^+/-/-^; V600E: A2058, A375, MCF10A^BRAF V600E^, WM3482, SK-MEL-5 and RKO. Each dot represents one cell line and the data are presented as mean ± SD. *p<0.05. **(P)** Representative DNQ-induced dose-dependent growth inhibition of CHL-1 and WM3482 cell lines. The experiments were repeated independently three times. The data are presented as mean ± SEM from triplicates from a representative experiment. **(Q)** Sequential combination effect of DNQ and vemurafenib in growth inhibition of WM3482 cell lines carrying BRAF V600E mutation. DNQ-induced dose-dependent growth inhibition of WM3482 cell viability were tested in three conditions: (1) DNQ alone, (2) pretreatment with 100nM vemurafenib for 24h followed by DNQ for 3 days (vemurafenib (1st) + DNQ (2nd)), and (3) pretreatment with DNQ for 24h followed by 100nM vemurafenib for 3 days (DNQ (1st) + vemurafenib (2nd)). The experiments were repeated independently three times. Data are presented as mean ± SEM from triplicates from a representative experiment. **(R)** AUC analysis of the combination effect of DNQ and vemurafenib in three melanoma cell lines, A2058, SK-MEL-5, and WM3482, with BRAF^V600E^ mutation. The experiments were repeated independently three times. Data are presented as mean of triplicates from a representative experiment. Each dot represents a cell line and the lines indicate mean ± SD. *p<0.05 from paired t-test. **(S)** Proposed working model of BRAF^V600E^/KEAP1 neoPPI in re-wiring KEAP1/NRF2/NQO1 ROS pathway and generating vulnerability to NQO1 substrate.

To examine the significance of the KEAP1 sequestration, we analyzed the CRISPR-Cas9 gene essentiality data from the DepMap (Meyers et al., 2017). BRAF^V600E^ mutations render cancer cells dependent on KEAP1 for cell survival with increased sensitivity to KEAP1 knockout (Fig. 6L, Table S10). Thus, it is possible that BRAF^V600E^ cells might inherit intrinsic sensitivity to KEAP1-controlled pathways.

To search for potential therapeutic agents to exploit BRAF^V600E^-induced vulnerability, we utilized a pair of genetically engineered isogenic MCF10A cell lines with V600E or WT in BRAF for chemical screening. Parallel cell viability screening with a library of bioactive compounds revealed a class of quinone compounds with selective growth inhibition of V600E cells (Fig. 6M). This result corroborated the observations from our pathway analysis that connected the BRAF^V600E^/KEAP1 interaction with enhanced activation of NQO1, a 2-electron reductase governing quinone metabolism (Dinkova-Kostova and Talalay, 2010). Thus, it is possible that the acquisition of the V600E mutation might lead to elevated NQO1 function with the consequential enhanced sensitivity to toxic NQO1 substrates. To test this contention, we utilized deoxynyboquinone (DNQ) (Fig. 6N), a potent and specific NQO1 substrate, to examine its effect on cell survival of BRAF^V600E^ cells (Parkinson et al., 2013). A panel of melanoma cell lines with differential BRAF genetic mutations was selected and tested for their sensitivity to DNQ in a dose-dependent cell viability assay. When treated with DNQ, BRAF^V600E^-carrying cells showed increased sensitivity over WT cells to DNQ with reduced AUC (Fig. 6O). Differential response curves were observed showing significantly enhanced sensitivity of the WM3482 cell line with BRAF^V600E^ compared to that of the CHL cell line with BRAF WT to DNQ treatment (Fig. 6P). Thus, BRAF^V600E^ in melanoma cancer cells may trigger enhanced sensitivity to NRF2-NQO1-modulated cytotoxic quinone derivatives.

Given that BRAF^V600E^/KEAP1 neoPPI can be regulated by BRAF^V600E^ inhibitor vemurafenib (Fig. 5F and 6I), we determined whether vemurafenib could synergize with DNQ in inhibiting the growth of V600E mutant cancer cells. Using the BRAF^V600E^ mutant WM3482 cell line as a model system, treatment with vemurafenib followed by DNQ showed a slightly decrease of DNQ’s potency compared to DNQ treatment alone, suggesting a negative effect of vemurafenib on DNQ (Fig. 6Q). However, when we reversed the order of treatment by pretreating cells with DNQ followed by vemurafenib, we observed a significant increase of DNQ’s potency (Fig. 6Q). Such potent sequential combination effects were also observed in multiple melanoma cell lines with the BRAF^V600E^ mutation (Fig. 6R). Thus, the neo-interaction of BRAF^V600E^ with KEAP1 not only re-wired the NRF2 regulated redox pathway, but also generated a cell state with enhanced sensitivity to cytotoxic NQO1 substrates. The uncovered BRAF^V600E^-variant directed sequential combination strategy supports future exploration of our demonstrated neoPPIs for mechanistic investigation and neoPPI-informed therapeutic strategies.

## DISCUSSION

Focusing on cancer driver mutations in this study for differential PPI discovery led to the emergence of a landscape where mutated residue-directed protein-protein interactions and pathways may rewire oncogenic programs to shape tumor phenotypes. A suite of neoPPI candidates have been revealed that suggests novel mechanisms for well-characterized somatic aberrations. The presence of neo-interactions stemming from genomic mutations seems widespread as they were found in all six genes tested, including both oncogenes and tumor suppressors. Also, neo-interactions were observed with different mutations in the same gene, or different alterations from the same residue, emphasizing the mutated residue-driven oncogenic programs. Such differential connectivity of diverse mutations may be important for the diverse clinical phenotypes and drug sensitivity displayed by different mutations in the same gene. The suggested molecular pathways mediated by these neoPPI candidates may provide unique opportunities to address variant-associated tumor heterogeneity and therapeutic responses.

Our results strongly support the notion that a mutated residue of an oncogenic driver may be displayed as a neo-epitope, or an induced neo-epitope, to change the composition of binding complexes and may enable the formation of neoPPIs for both oncogenes and tumor suppressors. For tumor suppressors, such a neo-epitope(s) may drive a “partner switching”, leading to not only the loss of tumor suppressor function, but also to a gain of oncogenic potential. For example, SPOP^F133L^ showed a reduced affinity for SRC and BRD4 while gaining an interaction with c-JUN, which may offer an oncogenic driver activity (Fig. 4E-H). Similarly, SMAD4^G386D^, while lost interaction with SMAD3, formed a neoPPI hub connecting with multiple oncogenic pathways, such as the Wnt signaling pathway (Fig. 4A-D). These neoPPIs may suggest potential pathway crosstalk contributing to SMAD4^G386D^ driven tumorigenesis and cancer progression. Such a driver mutation-based “partner switching” model may help address a major challenge in targeting the undrugged tumor suppressors, which is particularly significant because majority of cancer driver mutations occur in tumor suppressors. Those tumor suppressors without enzyme activities may also be amenable for PPI targeting. Thus, unexpected widespread gain-of-interaction neoPPIs with tumor suppressor mutations may provide actionable PPI targets for therapeutic intervention.

The uncovered oncogenic mutation-mediated Df-PPI and the prioritized neoPPI candidates present new hypothesis to address observed clinical complications and therapeutic responses of defined alterations, even in the field of molecularly targeted therapy. For example, the oncogenic kinase inhibitors have transformed cancer patient care. However, the available therapeutics that modulate the catalytic activity of the targeted kinases frequently result in limited clinical outcome, presenting a daunting challenge to understand the mutation-driven tumorigenic mechanism. As an example, AKT1^E17K^ is identified in up to 8% of breast cancers (Carpten et al., 2007; Jung et al., 2016). This mutation has been shown to enhance the protein kinase activity of AKT1 (Carpten et al., 2007). In support of this notion, several known AKT1 substrates, such as RICTOR, EZH2, HIF1B and SIN1, were identified as neoPPI binding partners, showing increased interaction signal with AKT1^E17K^. However, the molecular basis of the limited response rate observed in AKT1 inhibitor clinical trials, and an explanation of the paradoxical cellular context dependency of AKT1^E17K^ on cell growth control in spite of the protein kinase activation remains to be established (Beaver et al., 2013; Carpten et al., 2007; Lauring et al., 2010). The widespread AKT1^E17K^ induced neoPPIs suggest potential molecular connectivity underlying the observed context dependency and could inform potential drug combinations to improve the activity of AKT inhibitors. BRAF^V600E^ is another example as a clinically actionable target with FDA-approved kinase inhibitors. Despite the improved clinical benefit, primary and acquired resistance to BRAF inhibitors is observed frequently and develops almost inevitably (Hatzivassiliou et al., 2010; Sullivan and Flaherty, 2013; Sun et al., 2014), urging the interrogation of the molecular basis of BRAF^V600E^-driven oncogenic programs (Hatzivassiliou et al., 2010; Poulikakos et al., 2011; Tsoi et al., 2018; Vido et al., 2018). From qHT-dS, we identified a panel of BRAF^V600E^ neoPPI binding partners that inform the potential molecular connectivity between BRAF^V600E^ and various oncogenic pathways. The uncovered BRAF^V600E^/ACO1 neoPPI and the resulting ferritin upregulation raise the possibility of a BRAF mutation-regulated oncogenic program that is associated with perturbation of iron homeostasis as those in the ferroptosis sensitive melanoma cells (Tsoi et al., 2018). The BRAF^V600E^/KEAP1 interaction also informs ROS homeostasis pathway reprogramming (Fig. 4M-P).

Such mechanistic interpretations require in-depth neoPPI characterization and functional analysis. We presented the BRAF^V600E^/KEAP1 interaction as a case study to advance our neoPPI candidate studies. The BRAF^V600E^ mutation enables the binding of KEAP1, sequestration of KEAP1 away from NRF2, and the activation of NRF2-NQO1 redox signaling. Extensive molecular evaluations with a panel of biochemicaland cell biology approaches confirmed this neo-interaction *in vitro* in a direct manner and in cells under physiological conditions. The dynamic interaction complex among BRAF^V600E^, KEAP1, and NRF2 demonstrates a potential pathway by which BRAF^V600E^ contributes to tumorigenesis, and presents a promising molecular target for therapeutic intervention (Fig. 6S). Upon neoPPI validation, as a proof of concept, we examined the effect of the BRAF^V600E^/KEAP1 neoPPI on the NRF2 ROS pathway re-wiring and identified a potential vulnerability due to the upregulated NRF2-NQO1 pathway. Indeed, leveraging the NQO1 activated state for enhanced production of the cytotoxic DNQ metabolite led to the highly synergistic effect of a sequential combination approach with DNQ followed by vemurafenib. Such therapeutic insights informed by the discovery of the BRAF^V600E^/KEAP1 neoPPI exemplifies the utility of neoPPIs as a valuable resource for revealing hidden pathway dependencies and vulnerabilities.

Prevalent lineage diversities of recurrent mutations are observed in clinical tumor samples (Chang et al., 2016). In this study, we mapped the neoPPI profiles for several driver mutations found in diverse lineages, such as AKT1^E17K^ in breast cancer, BRAF^V600E^ in melanoma and thyroid cancers, and SPOP^F133L^ in prostate cancer. The recurrent neoPPI binding partners converge in these lineage-specific driver mutations onto several cancer-associated pathways, such as cell proliferation, cell cycle regulation and immune response (Fig. 3). Such pathway convergence suggests a potential common dependency of various tumor types on these reprogrammed signaling circuitries.

The Df-PPI mapping also links multiple mutations to the common cancer intrinsic JAK-STAT immune response pathway that has been proposed to mediate immune surveillance and checkpoint blockade therapy (Benci et al., 2016; Gao et al., 2016; Patel et al., 2017; Zaretsky et al., 2016). Mutations or loss of JAK-STAT pathway genes in cancer cells induce impaired immune response signaling, leading to primary or acquired immune resistance phenotypes (Benci et al., 2016; Gao et al., 2016; Patel et al., 2017; Zaretsky et al., 2016). Our study revealed multiple JAK-STAT pathway genes as recurrent neoPPI binding partners for mutations in multiple genes (Fig. 3). These JAK-STAT pathway associated neoPPIs provide promising molecular connectivity for further exploration of the potential functional significance of the JAK-STAT immune response pathway in mediating oncogenic mutations-induced immune escape and therapy resistance.

Our systematic discovery of driver mutant-induced Df-PPI and neoPPI candidates is enabled by the BRET^n^ technology-based qHT-dS platform that permits the identification of Df-PPIs at single amino acid resolution. The qHT-dS platform featuring live-cell PPI detection and streamlined protein expression monitoring and normalization provides a robust approach for differential PPI mapping in physiologically relevant environment, complementing the current computational, AP-MS and Y2H technologies (Cheng et al., 2021; Do et al., 2012; Sahni et al., 2015). The quantitative and comparative nature of the screening system coupled with the CARINA algorithm significantly enhances the data quality for Df-PPI discovery. Indeed, a significantly higher fraction of known WT PPIs was uncovered with the qHT-dS approach compared to the single-point analysis of the same dataset (Fig. 2C). Even though our primary purpose is to identify Df-PPIs, the comparison to human interactome datasets, such as BioGRID and BioPlex v2, showed that the qHT-dS PPI profiling has expanded the connectivity within the OncoPPi space by revealing new interactions and confirming previously reported PPIs (Oughtred et al., 2018; Rolland et al., 2014).

Together, our results support the concept of mutation-driven oncogenic pathway rewiring through mutant allele-enabled PPIs. The experimentally validated neoPPI candidates provide a mechanistic basis to generate testable hypothesis to further examine their functional significance. Such mutation-dictated PPIs may not only enhance our understanding of molecular reprograming in cancer, but may also reveal new ways to target tumor variant-mediated mechanisms for therapeutic intervention, which may be applicable to genomic alterations in both oncogenes and tumor suppressors. The presented Df-PPI dataset and the neoPPI candidates from our quantitative screening platform, bioinformatics annotation, and confirmative studies offer the scientific community a valuable resource for variant-directed molecular interaction studies to accelerate the precision medicine approach.

## ACKNOWLEDGEMENTS

We thank Dr. Paul J. Hergenrother for the DNQ compound, Dr. Josh Lauring for the isogenic MCF7 AKT1 WT and E17K cell lines, Dr. Donna Zhang for the pET15b-KEAP1 KELCH vector, RIKEN Brain Science Institute for the Venus plasmid, and Fu lab members for helpful comments and experimental assistance. The results published here are in part based upon data generated by the TCGA Research Network: http://cancergenome.nih.gov/. This work was supported by NCI’s Cancer Target Discovery and Development (CTD^2^) Network (U01CA217875 to HF; U01CA217842 to GBM). The data were deposited to the NCI-CTD^2^ Data Portal (https://ocg.cancer.gov/programs/ctd2/data-portal) and the CTD^2^ Dashboard (https://ctd2-dashboard.nci.nih.gov/dashboard/). This research was also supported in part by the NCI Emory Lung Cancer SPORE (SR, HF; P50CA217691) Career Enhancement Program (XM and AAI, P50CA217691), the Emory Chemical Biology Discovery Center, Winship Cancer Institute (NIH 5P30CA138292), Winship Cancer Institute #IRG-17-181-06 from the American Cancer Society (A.A.I.). Emory initiative on Biological Discovery through Chemical Innovation (A.A.I), the Imagine, Innovate and Impact (I3) Funds from the Emory School of Medicine and through the Georgia CTSA NIH award (UL1-TR002378), and a kind gift from the Miriam and Sheldon Adelson Medical Research Foundation to GBM. The authors have no conflicting financial interests. The authors plan on making the resource widely available to the academic community through CTD^2^ network and the Emory PPI portal.

## AUTHOR CONTRIBUTIONS

X.M., Q.N., A.A.I., Y.T.H., C.T., X.Y., C.S., S.D., D.C., D.F., and Y.D. conducted the experiments; X.M., Q.N., A.A.I., A.W., M.A.R., L.A.D.C., C.S.M., W.Z., T.O., S.L., F.R.K., Y.D., S.S.R., G.D.M, and H.F. participated in data analysis and interpretation; X.M., Q.N., A.A.I., G.B.M, F.R.K., Y.D. and H.F. participated in discussion and manuscript preparation; X.M., Q.N., D.C., A.A.I., Y.D. and H.F. designed the experiments and wrote the paper; and all were involved in manuscript editing.

## STAR METHODS

### RESOURCE AVAILABILITY

#### Lead Contact

Further information and requests for resources and reagents should be directed to and will be fulfilled by the Lead Contact, Haian Fu.

#### Materials Availability

Plasmids generated in this study are available upon request to the Lead Contact. Other materials are available through commercial sources (see Key Resource Table)

#### Data and Code Availability

Analyzed small molecule screening data sets are available through CTD^2^ data portal (https://ocg.cancer.gov/programs/ctd2/data-portal).

### METHOD DETAILS

#### OncoPPi v2 library of cancer-associated genes

Genes collected for the current studies are listed in Supplementary Tables S1-2. The OncoPPi v2 library includes all genes from the OncoPPi v1 gene set of lung cancer associated genes. To expand the OncoPPi library to other cancer types, we have identified the major tumor driver and tumor suppressor genes determined in large-scale cancer genomics studies, such as The Cancer Genome Atlas (TCGA) (https://www.cancer.gov/tcga). Based on the analysis of cancer-associated pathways defined in MSigDB, Reactome, and KEGG databases, we also included proteins involved in the regulation of oncogenic programs. All genes included in OncoPPi v2 library were annotated with the frequency of genomic alterations observed in the TCGA PanCancer patient cohorts available through the NCI Genomics Data Commons (GDC) Portal.

Each gene was subcloned into the indicated Gateway entry vector (Invitrogen). The integrity of the genes was confirmed by BsrGI restriction digestion and sequencing, generating the entry-vector library. Genes in the entry vector library were transferred using the Gateway recombination system to the destination expression vector to produce a Venus-flag-gene fusion for each gene, generating the OncoPPi v2 expression vector library. Site-directed mutagenesis was used to generate cancer driver mutation entry-vectors, which were fully sequenced and then transferred using Gateway recombination system to destination expression vector to produce a NLuc-HA-gene fusion for each cancer driver MUT and WT.

#### qHT-dS platform for Df-PPI screening

We used our previously established BRET^n^ technology in miniaturized uHTS 1536-well plate-based format to develop a qHT-dS platform to assess PPIs in live-cells for parallel comparative screening. NLuc- and Venus-fusion proteins allow streamlined monitoring of protein expressions simultaneously with BRET signal detection in a simple add-and-read mode. Briefly, HEK293T cells (1500 cells in 4 μl per well) were cultured in 1536-well plates at 37 °C before they were co-transfected in wells with Venus-tagged genes in combination with NLuc-tagged genes using Linear polyethylenimines (PEIs, Polysciences, 23966). Transfection was performed by adding 1 μl mixture of PEI (30 ng/μl) and DNA (10 ng/μl) to 4 μl cell culture, assisted by robotic operations with the Biomek NX^P^ Lab Automation Workstation (Beckman Coulter). BRET saturation assay was performed by titration of DNA amount to achieve various NLuc- and Venus-tagged gene combinations.

After incubation for 48 hours, Nano-Glo® luciferase substrate furimazine (Promega, N1120) was added to the cells directly. The donor luminescence signal at 460 nm and acceptor emission signal at 535 nm were measured immediately using an Envision Multilabel plate reader (PerkinElmer). The BRET^n^ signal is expressed as the ratio of light intensity measured at 535 nm over that at 460 nm. The specific BRET^n^ signal for the interaction of two proteins is expressed as net BRET^n^, which is defined as the difference in BRET signal with co-expression of two proteins and expression of the negative control NLuc-protein only.

The relative amount of NLuc-tagged protein expression was measured by the luminescence signal at 460 nm (L460) during the BRET^n^ signal measurement in 1536-well white plate (Corning, 3727); while the Venus acceptor protein expression was detected by the Venus fluorescence intensity (FI) with excitation at 480 nm and emission at 535 nm in 1536-well black clear-bottom plate (Corning, 3893). Cells were seeded and transfected side-by-side under the same conditions for the 1536-well white plate for BRET^n^ measurement and black plate for Venus FI measurement. The ratio of relative amount of acceptor over donor protein expression (Acceptor/Donor) was defined as Venus FI/L460. This intensity ratio should be proportional to the acceptor/donor ratio.

#### CARINA algorism for data analysis and statistics

To enable the rapid processing, quantification, and analysis of the qHT-dS data, we have developed the Comparative Analysis of Rewired INterActions (CARINA) algorithm, implemented as a Python Jupyter notebook. As the input data, CARINA directly uses the sets of raw data files generated by the plate reader in 1536-well plate format, including the BRETn PPI signals and Venus fluorescence intensity readouts. Based on the pre-designed plate templates, CARINA automatically recognizes the signals that correspond to PPI and empty vector controls for a particular bait mutated protein. For each individual PPI, CARINA computes three BRET^n^ saturation curves, including one curve for PPI and two curves for empty NLuc (ctrl1) and Venus (ctrl2) controls, with each in four replicates. The saturation curves are based on the equation

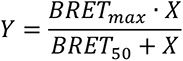

where Y is net BRET^n^, and X is Acceptor/Donor. The negative X and Y values were excluded from the analysis. For quantitative analysis, the area under the curve (AUC) was computed as a measurement of BRET^n^ signal, as AUC integrates both the amplitude and shape of the saturation curve (Mo and Fu, 2016; Mo et al., 2016). The fold-over-control (FOC) was calculated as AUC_PPI_/AUC_max(ctrl1, ctrl2)_ to express the difference between PPI and empty vector controls, and statistical significance (P_FOC_) was calculated using Student’s t-test to estimate the likelihood that AUC_PPI_ is different from the AUC_ctrl1&ctrl2_.

To express the differences between the WT and MUT PPI profiles, CARINA calculates the differential score (DS) and p-value of DS (P_DS_) that estimate the likelihood that FOC_WT_ is significantly different from FOC_MUT_. When WT PPI is negative and MUT PPI is positive, DS=FOC_MUT_ and P_DS_=P_FOC_(MUT). When WT PPI is positive and MUT PPI is negative, DS=1/FOC_WT_ and P_DS_=P_FOC_(WT). When both WT and MUT PPIs are positive, DS will be defined as FOC_MUT_/FOC_WT_ and P_DS_ will be calculated using Student’s t-test of FOC_WT_ and FOC_MUT_. The Benjamini-Hochberg false discovery rate (FDR)-adjusted P_DS_ are calculated based on the Student’s t-test of AUC_WT_ and AUC_MUT_ to prioritize neoPPI by considering the potential artifacts from previous statistical cut-offs. Python protocols and Matlab scripts and functions used in data processing and analysis are available on request.

#### Gene pathway enrichment analysis

The enrichment of the OncoPPi v2 library in cancer-associated genes has been done in python using the Enricher API (Xie et al., 2021) and based on the KEGG 2021 Human and MSigDB Hallmark 2020 gene sets.

For pathway analysis of Df-PPIs, we used a list of 125 genes that have ≥3 interaction mutation alleles. In this list, we looked for pathway category over-representation using Ingenuity Pathway Analysis (QIAGEN, Redwood City, http://www.qiagen.com/ingenuity). The analysis criteria were set as follows: (1) querying for molecules with Ingenuity Knowledge Base as a reference set; (2) restricted to human species; and (3) experimentally observed findings as a confidence level. Fisher’s exact test (p-value < 0.05) was used to compute significance for over-representation of genes in a particular pathway or biological process.

#### Molecular biology techniques and cell culture conditions

Individual cloning vectors, site-directed mutagenesis and truncation mutants were made and verified by following product manual and standard molecular biology protocols. Human cancer cell lines (CHL-1, HMCB, A2058, and SK-MEL-5 from ATCC, Manassas, VA; WM3311, and WM3482 from Rockland Immunochemicals Inc., Limerick, PA; parental MCF10A, isogenic BRAF^V600E^ knockin MCF10A, parental RKO, and isogenic BRAF V600E mutant alleles knockout RKO from Horizon Discovery, Saint Louis, MO) were cultured in cell culture medias suggested by manufacturer. HEK293T cells (ATCC, Manassas, VA) were maintained in regular Dulbecco’s Modified Eagle’s Medium (DMEM, Corning, 10013CV), supplemented with 10% fetal bovine serum (FBS, Sigma, F0926) and 1x penicillin/streptomycin solution (Corning, 3001CI), or in phenol red free DMEM (HyClone, SH30284.01) throughout the screening. Cells were incubated at 37 °C in humidified conditions with 5% CO_2_. For drug treatment for functional studies, cells were treated with either vehicle or chemicals at indicated concentration.

#### Affinity pull-down and co-immunoprecipitation assays

For GST-affinity pull-down assays, cells were lysed in NP-40 buffer (1% nonident P-40, 20 mM Tris ((PH 8.0), 137 mM NaCl, 5% glycerol, protease inhibitor (Sigma, P8340) and phosphatase inhibitor (Sigma, P5726 and P0044)) and incubated with glutathione-conjugated beads (GE,17075605) for 2 hours at 4 °C. Beads were washed three times with 1% NP-40 buffer and eluted by boiling in Laemmli sample buffer (Bio-Rad, 1610737). For immunoprecipitation (IP), cell lysates were collected, quantified, and were mixed with respective antibodies. For each co-IP, lysates containing ∼1.5 mg of total proteins were used and the antibody/lysate mixtures were incubated overnight at 4 °C. Then protein A/D agarose beads (Santa Cruz, sc-2003) were added to the mixture followed by incubation at 4 °C for another 4 hours. Beads were washed four times with lysis buffer, and proteins were eluted with SDS-PAGE sample buffer and analyzed with indicated antibodies. The following primary antibodies were used for Western blotting: rabbit anti-GST (Sigma, A7340); mouse anti-Flag (Sigma, A8592); rabbit anti-ERK1/2 (Cell Signaling, 9102); rabbit anti-phospho-ERK1/2 (Cell Signaling, 4370); rabbit anti-NRF2 (Abcam, ab31163); rabbit anti-KEAP1 (Cell Signaling, 4678); mouse anti-β-Actin (Sigma, A5441); mouse anti-NQO1 (Santa Cruz, sc-32793); mouse anti-BRAF (Santa Cruz, sc-5284).

#### Venus protein-fragment complementation assay (PCA)

The Venus-PCA was performed as described previously (Pham, 2015). Briefly, the cells were co-transfected with BRAF WT vs V600E and KEAP1 conjugated to C-terminal or N-terminal fragments of Venus. The cells transfected with empty vector as controls. Proteins were expressed for 48 hours, and the protein-protein interactions were monitored in live cells based on the fluorescence intensity of reconstituted Venus. The cell nuclei were stained with 5 μg/ml Hoechst 33342 (ThermoFisher, H3570) for 30 minutes at 4 °C. The cell images were taken with ImageXpress system at 447 nm (Hoechst nuclei stain) and 530 nm (Venus green fluorescence). The fluorescence intensity of reconstituted Venus was measured on the Envision spectrophotometer (Ex 485 nm and Em 535 nm, mirror 505 nm).

#### BioLayer Interferometry (BLI)

Direct binding studies were carried out using the Octet Red 384 system (Forte Bio), at 30 °C with shaking at 1,500 RPM in a 384-well plate containing 50 μl of the solution in each well. 1X BLI Kinetic Buffer (PBS, pH 7.4, 10 mg/ml bovine serum albumin (BSA) and 0.1% (v/v) Tween 20) was used throughout this study for protein dilution and washing steps. Ni-NTA biosensors (Sartorius, 18-5101) were balanced for the first baseline and then loaded with His-KEAP1 KELCH domain (25 μg/ml) followed by another wash for the second baseline. Kinetic analysis of the interaction was performed by dipping the sensors into the well containing GST-BRAF WT or V600E kinase domain (200 nM).

#### RNA isolation, cDNA synthesis and qPCR

Total RNA was isolated using the Direct-zol^TM^ RNA Mini-Prep kit (Zymo Research, R2071) following the manufacturer’s instruction. cDNA synthesis was performing with the SuperScript III First-Strand Synthesis System (Life Technologies, 18080-051). Primers for qPCR analysis of gene expression are listed in Supplementary Table (?). The expression of β-Actin and GAPDH were used as control for signal normalization.

#### Protein stability assays

HEK293T cells were grown in 24-well plates and transfected using FuGene (Promega, E2311) following the manufactureŕs instructions. After transfection, cells were incubated for 48 hours in DMEM media supplemented with 10% FBS, then were treated with 5 μg/mL cycloheximide (Cell Signaling, 2112) in DMEM media with 10% FBS. At the indicated times, 100 μl of 2X SDS-PAGE sample buffer was added and the cells were scraped from the wells, boiled for 5 minutes, then cell lysates were stored at -80 °C. After all lysates were collected, each sample was loaded onto a 10% SDS-PAGE gel and then analyzed by Western blotting with anti-NRF2 antibody to monitor NRF2 protein level. Protein expression was quantified from the Western blot using ImageJ software for analysis, NRF2 levels were normalized to β-Actin protein levels. Assays were performed three times.

#### Luciferase reporter assay

HEK293T cells were grown in 6-well plates and transfected using FuGene with Venus-flag-BRAF WT, or V600E, or Venus vector along with NRF2-ARE, or TCF/LEF, or AP-1 luciferase reporter plasmids (BPS Bioscience, 60514; Promega, E4661; Qiagen, CCS-011L). Renilla luciferase was included as an internal control. After transfection, cells were incubated for 48 hours in DMEM media supplemented with 10% FBS. Cells were harvested mechanically, centrifuged at 1,600 rpm for 2 minutes, then re-suspended in 300 μl of DMEM media. The cells were transferred to 384-well plate, and the reporter assay was performed using Dual-Glo luciferase kit (Promega, E2920) following the manufacturer’s instructions. Firefly luciferase expression was normalized to the internal control Renilla expression. Data were analyzed with Graphpad Prism software. Assays were performed three times.

#### HTS cell viability assay

Cells were seeded at 3000 cells/well in 50 μl media in a 384-well culture plate (Corning, Cat#3764) using a Multidrop™ Combi Reagent Dispenser (ThermoScientific) with the first column as a medium only control (blk). The next day test compounds (0.1 ul) were dispensed into wells in each plate using a Sciclone ALH 3000 liquid handler (PerkinElmer) from a compound stock plate to give the indicated final concentrations. Each sample was tested with 3 replicates. After 3 days of incubation, 10 μl CellTiter-Blue (Promega, G8081) was added to each well using the robotic liquid dispenser. The plate was incubated for 1-4 hours at 37 °C. The fluorescence intensity (FI) of each well was read using an EnVision Multilabel plate reader (Ex 545 nm, Em 615 nm, PerkinElmer). Percentage (%) of Control was calculated using the equation: (FI compound - FI Avg. Blk)/ (FI Avg. Neg. - FI Avg. Blk * 100).

## Figure legends for supplementary figures

Figure S1. qHT-dS platform design and illustration, related to Fig. 1B.

Figure S2. OncoPPi v2 library design, related to Fig. 1C.

Figure S3A. GST-pull down validation of SMAD4-G386D neoPPI, related to Fig. 4A.

Figure S3B. GST-pull down validation of SPOP-F133L neoPPI, related to Fig. 4E.

Figure S3C. GST-pull down validation of FBXW7 and SMARCA4 neoPPI.

Figure S3D. GST-pull down validation of AKT1-E17K neoPPI, related to Fig. 4I.

Figure S3E. GST-pull down validation of BRAF-V600E neoPPI, related to Fig. 4M.

Figure S4. Mutual exclusivity analysis of BRAF-V600E and its neoPPI binding partners, related to Fig. 4M.

Figure S5. Endogenous co-immunoprecipitation validation of BRAF-V600E/KEAP1 neoPPI in melanoma cell lines, related to Fig. 5C.

## Supplementary Tables

Table S1. OncoPPi v2 library gene ID and enrichment analysis. Five genes, including AXIN2, CDKN2A, NKX2-1, NSD3, and SRSF2, have two ORFs included.

Table S2. Summary of genetic alterations in OncoPPi v2 library genes.

Table S3. Raw data from qHT-dS and calculated X and Y for BRET saturation curve analysis.

Table S4. CARINA analysis results.

Table S5. Pathway enrichment analysis.

Table S6. Analysis of SMAD4^G386D^ neoPPI binding partners that co-localize with SMAD4.

Table S7. Analysis of AKT1 neoPPI binding partners that are known substrates, with putative AKT1 phosphorylation consensus motif RxRxxS/T, or co-localize with AKT1.

Table S8. Mutual exclusivity, co-localization, CRISPR, RNAi and drug sensitivity analysis of BRAF V600E and its neoPPI binding partners.

Table S9. Correlation analysis of BRAF genetic status with NQO1 mRNA expression from CCLE dataset.

Table S10. The essentiality of KEAP1 gene knockout in BRAF WT and V600E genes determined in CRISPR-Cas9 screen in 436 cancer cell lines.

